# Chromatin-enriched RNAs mark active and repressive cis-regulation: an analysis of nuclear RNA-seq

**DOI:** 10.1101/646950

**Authors:** Xiangying Sun, Zhezhen Wang, Carlos Perez-Cervantes, Alex Ruthenburg, Ivan Moskowitz, Michael Gribskov, Xinan Holly Yang

**Affiliations:** Department of Biological Sciences, Purdue University, West Lafayette, IN 47906, USA; Department of Pediatrics, The University of Chicago, Chicago, IL 60637, USA; Department of Molecular Genetics and Cell Biology, The University of Chicago; Department of Computer Science, Purdue University, West Lafayette, IN 47906, USA

**Keywords:** Computational genomics, enhancer, nuclear RNA-seq, long noncoding RNA, gene regulation

## Abstract

Long noncoding RNAs (**lncRNAs**) localize in the cell nucleus and influence gene expression through a variety of molecular mechanisms. RNA sequencing of two biochemical fractions of nuclei reveals a unique class of lncRNAs, termed chromatin-enriched nuclear RNAs (**cheRNAs**) that are tightly bound to chromatin and putatively function to cis-activate gene expression. Until now, a rigorous analytic pipeline for nuclear RNA-seq has been lacking. In this study, we survey four computational strategies for nuclear RNA-seq data analysis and show that a new pipeline, Tuxedo, outperforms other approaches. Tuxedo not only assembles a more complete transcriptome, but also identifies cheRNA with higher accuracy. We have used Tuxedo to analyze gold-standard K562 cell datasets and further characterize the genomic features of intergenic cheRNA (**icheRNA**) and their similarity to those of enhancer RNA (**eRNA**). Moreover, we quantify the transcriptional correlation of icheRNA and adjacent genes, and suggest that icheRNA may be the cis-acting transcriptional regulator that is more positively associated with neighboring gene expression than eRNA predicted by state-of-art method or CAGE signal. We also explore two novel genomic associations, suggesting cheRNA may have diverse functions. A possible new role of H3K9me3 modification coincident with icheRNA may be associated with active enhancer derived from ancient mobile elements, while a potential cis-repressive function of antisense cheRNA (**as-cheRNA**) is likely to be involved in transiently modulating cell type-specific cis-regulation.

**Author Summary:** Chromatin-enriched nuclear RNA (cheRNA) is a class of gene regulatory non-coding RNAs. CheRNA provides a powerful way to profile the nuclear transcriptional landscape, especially to profile the noncoding transcriptome. The computational framework presented here provides a reliable approach to identifying cheRNA, and for studying cell-type specific gene regulation. We found that intergenic cheRNA, including intergenic cheRNA with high levels of H3K9me3 (a mark associated with closed/repressed chromatin), may act as a transcriptional activator. In contrast, antisense cheRNA, which originates from the complementary strand of the protein-coding gene, may interact with diverse chromatin modulators to repress local transcription. With our new pipeline, one future challenge will be refining the functional mechanisms of these noncoding RNA classes through exploring their regulatory roles, which are involved in diverse molecular and cellular processes in human and other organisms.

## 1. Introduction

Long noncoding RNA (**lncRNA**) is enriched not only in the cell nucleus, but also within the chromatin-associated fraction [1]. Many nuclear lncRNAs affect coding gene expression and chromatin organization, and are important in diverse biological processes [2, 3]. Among these nuclear lncRNAs, chromatin-associated lncRNA (**cheRNA**) processes gene-regulatory roles [4–6]. In our recent studies, cheRNA promotes essential gene-enhancer contacts and is associated with transcript factor-dependent lncRNA [5, 7]. However, a robust analytic pipeline for cheRNA as a group of functional nuclear RNAs has not been performed.

Bioinformatic efforts on nuclear RNA-seq are required because the distinctions between mRNA-seq and nuclear RNA-seq are substantial. The differences on library construction have significant consequences for the interpretation and analysis of the sequencing data [8]. For instance, sequencing of polyadenylated (polyA+) RNA may miss transcripts that are not usually polyadenylated, which includes many lncRNAs. Total-RNA sequencing detects a higher proportion of lncRNA, but is more expensive and less efficient in quantifying coding-gene expression [9].

In this study, we compare one published and three new analytic pipelines for nuclear RNA-seq data analysis. A newly developed pipeline, Tuxedo, outperforms the other pipelines with respect to transcriptome completeness, accuracy of cheRNA identification, and enrichment of enhancer-hallmarks at cheRNA gene regions. Using the Tuxedo pipeline, we identify cheRNAs genome-wide. Based on the GENCODE annotation (v25) [10], we verify that cheRNA provides a new way to annotate putative enhancer RNA (**eRNA**).

We explore the functions of two subsets of cheRNA with potential regulatory functions but new genomic markers: intergenic cheRNA transcribed from genes in condensed chromatin (marked by H3K9me3), and cheRNA transcripts antisense to coding genes. H3K9me3 is primarily recognized for transcript repression. Surprisingly, our data indicate that cheRNA-marked eRNAs are located within regions of high H3K9me3. Given H3K9me3 is associated with mobile elements and their transcription [11, 12], our data agrees with the theory that enhancers are co-opted from ancient mobile elements [13]. CheRNA may have subgroups, notably is the one antisense to coding genes distinctly showing negative correlations in transcription level with their corresponding sense mRNA. Our approach affords a straightforward approach to identifying and predicting novel regulatory lncRNA for future mechanistic evaluation.

## 2. Results

### 2.1. Nuclear RNA-seq requires rigorous computational strategies

Nuclear RNA-seq library construction differs from other RNA-seq protocols (Fig 1a). For example, the numbers of detected transcripts differ when nuclear or total RNA is sequenced, with 30.0% (7.0 k out of 23.3 k) of the transcripts detected only by total RNA sequencing, and 15.9% only by nuclear (Fig 1b). This difference is unlikely to be simply due to sequencing depth -- the median depth was 49M for four pooled total RNA samples and 33M for 22 nuclear RNA samples; the latter includes the 9 CPE and 9 SNE samples reanalyzed in this study (**Table S1**). Markers of transcriptional regulation including RNA polymerase II (**Pol II**) sites, transcription factor binding sites, cis-regulatory RNA structures, histone deacetylase, and histone enhancer hallmarks are common in the DNA corresponding to the 3700 RNAs detectable only by nuclear RNA-seq (**Fig S1, Table S2**). This observation agrees with previous suggestions that nuclear-retained lncRNA may interact with chromatin regulatory proteins and recruit them to cis-regulatory elements in order to regulate gene expression [1, 3]. Therefore, nuclear RNA-seq requires rigorous and effective pipelines different from the conventional pipelines used for total RNA-seq datasets.

**Fig 1.**
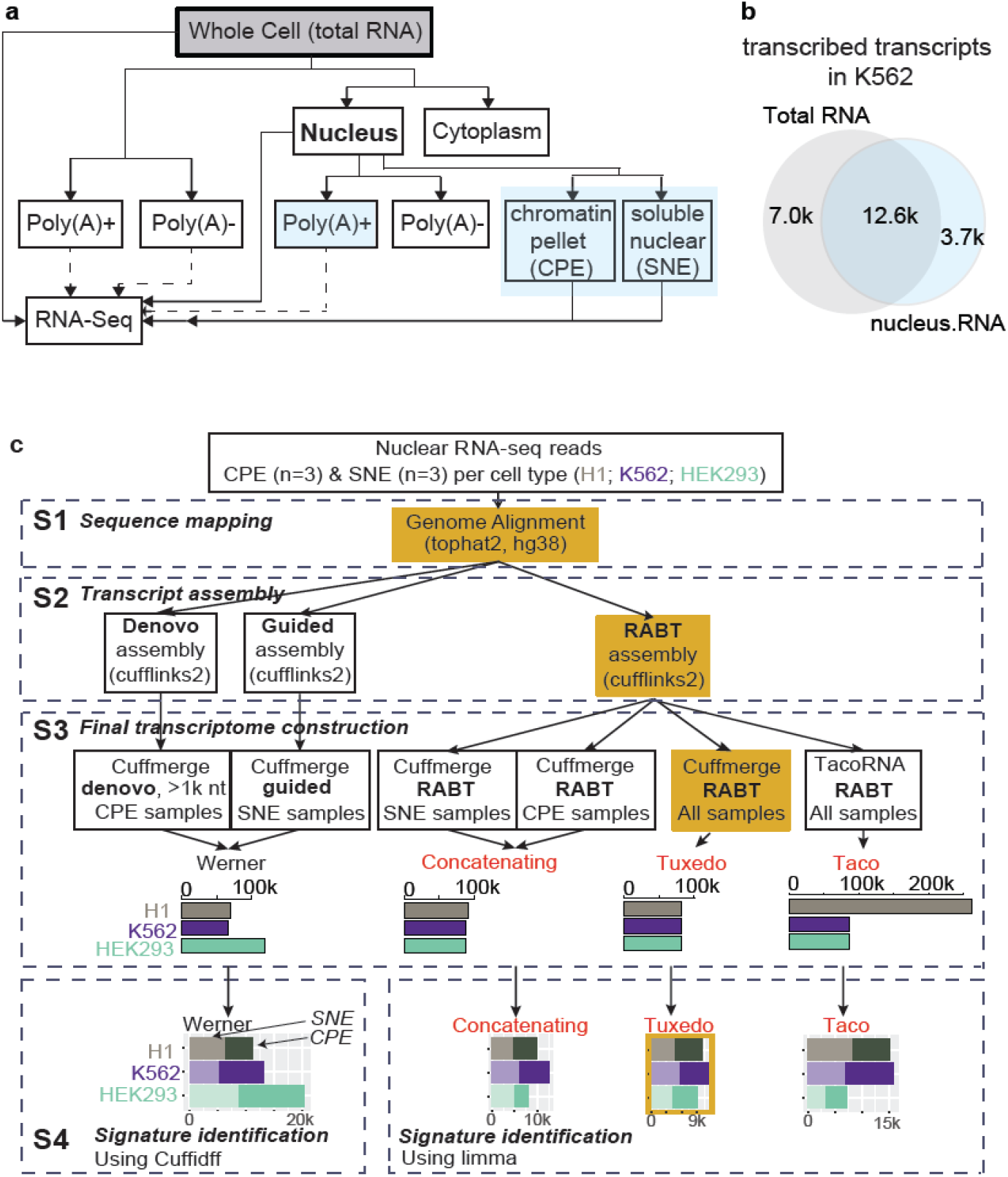
Nuclear RNA-seq sheds new insights into cis-regulatory elements. **a**, Diverse RNA-seq library strategies from parallel samples. Solid lines are the sequencing libraries (in category) analyzed in this study, and dashed lines are other available libraries. Blue color indicates the nuclear RNA-seq strategies. **b**, Venn diagram compares the number of predicted transcripts in two pooled RNA-seq transcriptomes (K562 cells). One is the total RNA-seq transcripts that were expressed with ENCODE transcript quantification value>0 in both replicates, in at least one of four collected samples, and the other is the three types of nuclear RNA-seq transcripts (blue boxes in panel a, pooled from 22 samples (see **Table S1**). The latter includes either all detectable transcripts (those with non-NA values in the downloaded data) in both its replicates, or Tuxedo-assembled expressed transcripts with CPM ≥ 1. **c**, Workflow of the four nuclear RNA-Seq analytic pipelines with four major analytic steps: **S1**, Sequence mapping. **S2**, Transcript assembly. **S3**, Final transcriptome construction. Bar plot represents the number of “expressed” transcripts (CPM ≥ 1 in at least two samples); color indicates the assembly result in different cell line. **S4**, Signature identification. Stacked bar plot represents the number of RNA with different abundance in the CPE (darker color) or SNE samples (lighter color).

After the isolation of RNA and generation of sequencing libraries, a typical RNA-seq analytic workflow involves sequencing hundreds of millions of reads, alignment of reads against a reference genome or transcriptome, and downstream statistical analysis of expression. In the original method developed by Werner *et al.* [4, 5], which we refer to as Werner, cheRNA was identified in four steps (**S Method**). This Werner pipeline has three important biases: 1) Werner overestimates the proportion of *de novo* transcripts originating from CPE because it applies reference-guided *de novo* assembly (which can discover novel transcripts) to CPE but not to SNE fractions. 2) Werner removes transcripts shorter than 1000 bases from the analysis. LncRNA transcripts are typically shorter than (median length 592 bases) protein-coding transcripts (median 2.4k bases), and 33% of GENCODE-annotated noncoding RNA is shorter than 1000 bases long [14]. Removing transcripts shorter than 1,000 bases from the CPE assembly leads to significant under-detection of lncRNA. 3) In the differential expression analysis, Cuffdiff was used in Werner. However, Cuffdiff cannot do a two-group test on RNAs that have high expression levels. For example, noncoding RNA *XIST*, which is a canonical cheRNA, was categorized as “HiDATA” and excluded from differential expression analysis by Cuffdiff. In addition, it has been shown that discarding genes that are not expressed at a biologically meaningful level in any condition (prefiltering) can increase the power for detecting differential gene expression [15], but Werner doesn’t include a prefiltering step in the differential expression analysis.

### 2.2. Tuxedo outperforms three existing and alternative analytic pipelines

#### Tuxedo builds a complete transcriptome for active transcripts

We developed three new pipelines (referred to here as Tuxedo, Concatenating, and Taco) to analyze these datasets; each with four major analytic steps (Fig 1c). Detailed discussion about the four nuclear RNA-seq analytic pipelines in theory is given in the **S Method**. The Tuxedo pipeline makes three key computational improvements: 1) Tuxedo assembles the complete transcriptome in an unbiased way, covering both highly-expressed transcripts and lncRNA shorter than 1,000 bases. 2) Tuxedo employs an empirical threshold to distinguish between low but informative lncRNA transcription and noise. And 3) for the first time, we report a correlation between the expression of icheRNA and adjacent genes, facilitating the prioritization of potentially cis-acting cheRNA for further experimental evaluation. The strategies used in the Tuxedo pipeline are not restricted to cheRNA identification, and could be beneficial to nuclear RNA-seq data analyses testing broader biological hypotheses, such as to the relationship between enhancer marked (e.g., **Fig S1**) and differentially expressed nuclear RNAs.

Given that lowly expressed transcripts are likely to be experimental noise [16], we adopted Tuxedo to identify lncRNAs that may expressed lowly. Unlike methods applied to coding gene profiles, in which one can define an expression cutoff for active promoters, we made an empirical decision to define predicted transcripts with CPM (counts per million) ≥ 1 as ‘expressed’ for downstream analysis. This filter resulted in an approximately log-normal distribution of expression levels and about 14 k measured transcripts per sample (**Fig S2a**), ensuring the appropriateness of model-based differential expressional analyses such as Limma [17].

To evaluate the transcriptomes assembled by the four pipelines, we used the assembly result of K562 cell data as a reference. We first compared the completeness of the transcriptomes. Transcriptomes assembled by the Tuxedo and Concatenating assemblies are very concordant. 99.8% of transcripts are the same. 84.4% of transcripts assembled by both Tuxedo and Concatenating are also assembled by Werner. This number decreases to 27.0% for Taco (Fig 2a). To determine the reasons for assembly inconsistency, we compared the assembly results for annotated transcripts (Fig 2b) and unannotated transcripts (Fig 2c) separately. The annotated transcripts assembled by Tuxedo, Concatenating and Werner pipelines are almost identical, and correspond to the set of annotated transcripts in GENCODE (v25). Taco only assembled 26.2% of annotated transcripts. This is because the Taco pipeline uses TACO instead of Cufflinks as the assembly tool. TACO only includes transcripts that have significant expression, while Cufflinks keeps all annotated transcripts when building the transcriptome. By looking at the expression level of transcripts, we confirmed that the 40.7 k annotated transcripts that are omitted by Taco are transcripts with low expression levels in K562. Among the unannotated transcripts assembled by Tuxedo and Concatenating, 96.8 % are the same. Moreover, Tuxedo and Concatenating assembled more transcripts than Werner and Taco. The length distribution of transcripts assembled by Tuxedo and Concatenating, but not by Werner, shows that the majority of such transcripts are shorter than 1000 bases (Fig 2d), which is caused by removing transcripts shorter than 1000 bases from CPE samples in Werner. Taco assembled the smallest number of unannotated transcripts. The unannotated transcripts omitted by Taco also have low expression. Even though these transcripts were lowly expressed in samples, it is still necessary to keep them in the assembled transcriptome to accurately estimate gene expression levels.

**Fig 2.**
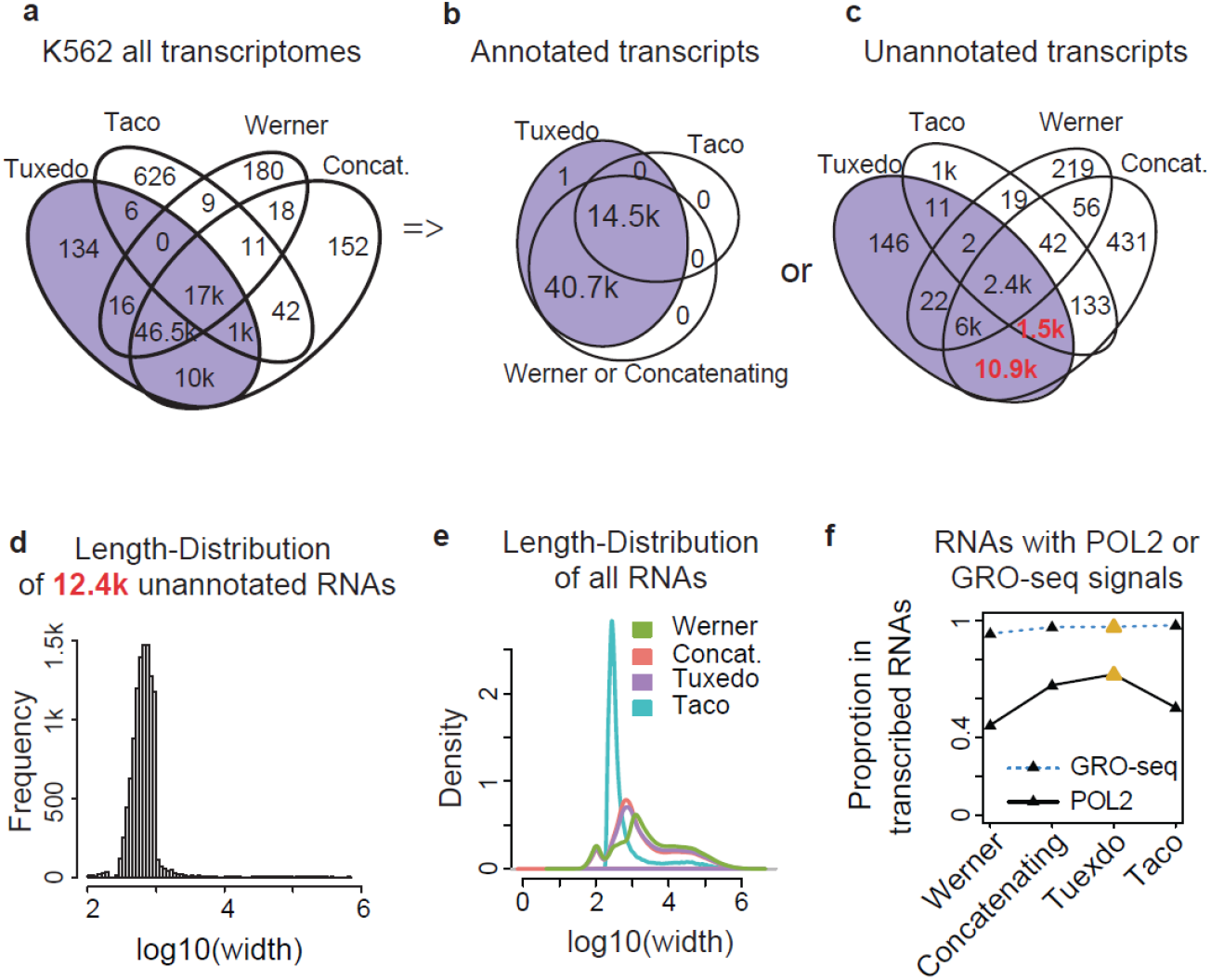
Tuxedo assembles a complete high quality transcriptome. **a**, Coordinate-overlap for all transcripts in predicted RNA classes in the K562 cell. Numbers are counted by the R package ChIPpeakAnno with the “findOverlapsOfPeaks” function. RNAs with an overlap of 1 base or more are considered to be overlapped. If multiple transcripts overlap in several groups, the minimal number of transcripts in any group is shown. **b**, Coordinate-overlaps for annotated or **c**, unannotated RNA, being respectively constructed by 4 pipelines. **d**, Length distribution of unannotated RNA predicted by Tuxedo and Concatenating but not by Werner or Taco. **e**, Length distribution of all predicted RNA in the four pipelines. **f**, Proportion of expressed RNA (CPM ≥ 1) assembled in each pipeline that overlap (at least 1 base, same strand) by coordinate with any GRO-seq peak and POL II peak.

We next investigated the length of assembled transcripts (Fig 2e, **Fig S2b**). Approximately half of the assembled transcripts have lengths between 200-1000 bases and show similar log-normal distributions for Tuxedo and Concatenating. In contrast, transcripts assembled by Werner are generally longer (71% of the assembled transcripts are longer than 1000 bases), which is another indication of the incompleteness caused by removing short transcripts. Transcripts assembled by Taco are much shorter (83% of assembled transcripts are shorter than 1000 bases). The TACO assembler employs an algorithm based on change-point detection via binary segmentation to predict transcript structure [18]. This algorithm is more robust in the assembly of annotated and conserved transcript such as mRNA. However, when it is applied to assembly of noncoding RNA, the TACO assembler overestimates the degree of alternative splicing and results in a large number of truncated transcripts. This is incorrect since only a small fraction of lncRNA undergo splicing [19]. In summary, Tuxedo and Concatenating construct relatively complete and correctly structured transcriptomes for analysis.

Additionally, we compared the transcriptional activity of the expressed transcripts assembled by the four pipelines using two independent measurements: Pol II ChIP-seq and global run-on sequencing (**GRO-seq**) (**Table S1**). Expressed transcripts are defined as those having CPM ≥ 1. The expressed transcripts assembled by Tuxedo show the highest proportion of overlap with peaks representing both ongoing transcription by Pol II and peaks representing nascent transcription by GRO-seq (Fig 2f), demonstrating that expressed transcripts assembled by Tuxedo are more concordant with active transcription signal represented by other methods. In summary, we introduce the computational pipeline, Tuxedo, for analyzing nuclear RNA-seq data containing both high low expression lncRNA.

#### Tuxedo identifies cheRNAs precisely while recapturing three known genomic features of active enhancers

To evaluate the performance of the four pipelines in cheRNA identification, we used the set of transcripts identified by all methods as a proxy gold standard, and found Tuxedo and Concatenating outperformed Werner and Taco in the identification of both cheRNA and sneRNA (Fig 3a). To further check the accuracy we examined sixteen loci of known cheRNA, sneRNA, or chromatin-independent RNA (transcripts not significantly differentially expressed between CPE and SNE samples) that were previously validated in specific cell types [4, 5]. Tuxedo and Concatenating successfully confirmed the chromatin enrichment in all canonical cheRNAs and outperform Werner and Taco with overall positive predicted value (**ppv**) of 0.88 (Fig 3b, **Table S3**). This analysis, although possibly susceptible to threshold effects, makes up the shortage of lack of a true gold standard in the ROC-analysis. Both analyses suggest that Tuxedo and Concatenating are better than Werner and Taco.

**Fig 3.**
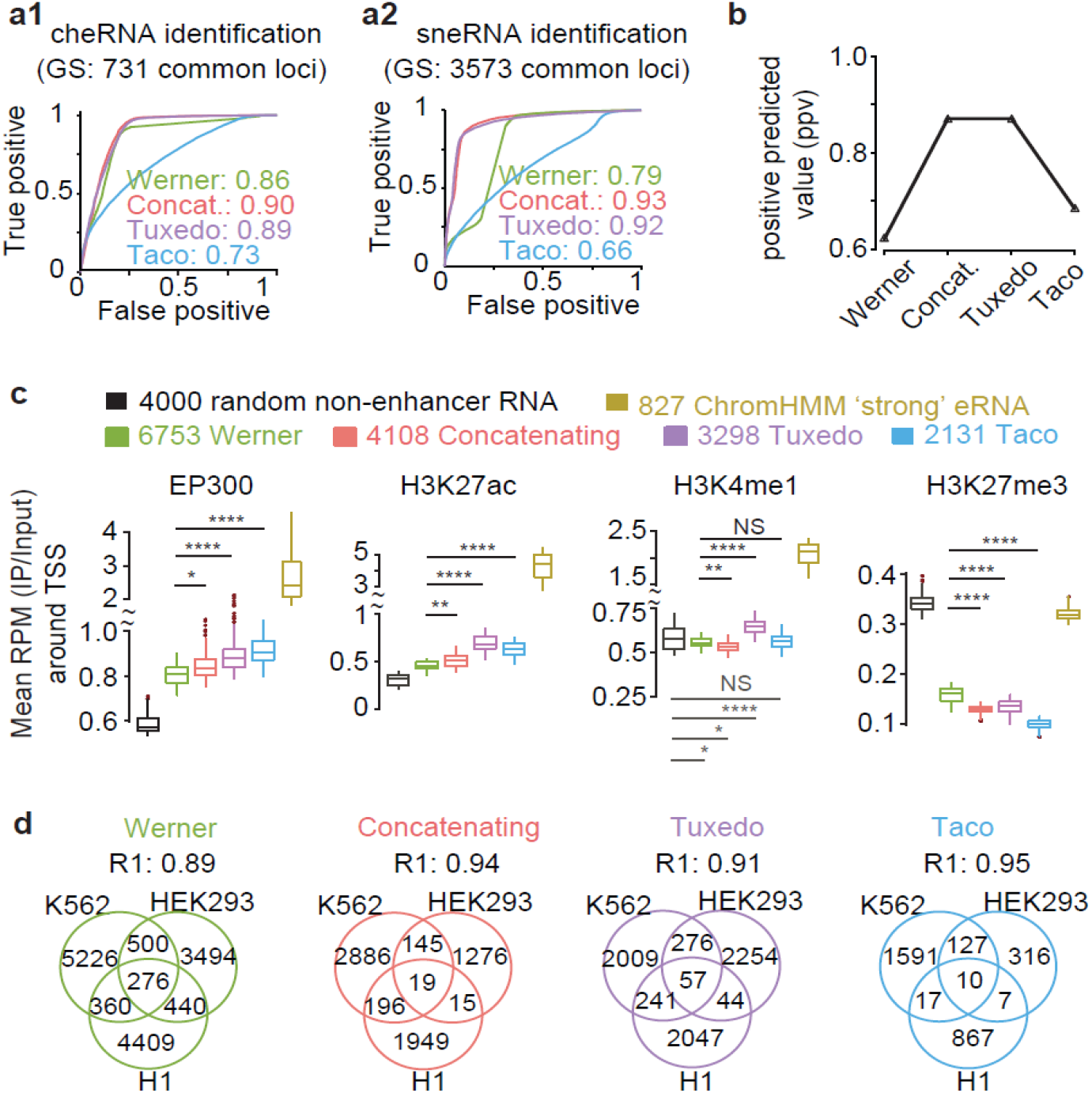
cheRNA prediction using the four pipelines in the K562 cell line. **a**, Receiver operating characteristic (ROC) curves of four pipelines identifying cheRNA (a1) and sneRNA (a2). The commonly-identified 731 cheRNA or 3573 sneRNA by all four pipelines are the proxy gold standard (GS) used here. **b**, Average positive predicted value (**ppv**) in identifying sixteen canonical cheRNA/sneRNA/intermediate RNA (RNA not differentially expressed between CPE and SNE) experimentally verified in previous studies for chromatin-enrichment or depletion, using the four pipelines, respectively. Further details about these 16 loci are given in **Table S3**. **c**, Average ChIP-seq read density around TSS (±1kb centered at TSS) of the indicated RNA classes in K562. Boxes span the lower to upper quartile boundaries, the median is indicated with solid line in each box. ChromHMM-predicted eRNA is defined as intergenic RNA overlapped (at least 1 base, same strand) with any ChromHMM predicted “strong enhancer” region and ChromHMM-predicted non-enhancer RNA is defined as transcribed RNA that have no overlap with any ChromHMM-predicted “strong” or “weak” enhancers. **d**, Fraction of cell-type-specific intergenic cheRNAs. R1 is the ratio of cell type specific RNAs versus all RNAs identified in each pipeline. Higher R1 value indicating more cell-type specific identification. Venn diagrams show the overlap of icheRNA identified in K562, HEK293 and H1-hESC cell lines by Werner (green), Concatenating (red), Tuxedo (purple) and Taco (blue). icheRNA identified by all the four pipelines except Werner (green) have high tissue-specificity (R1>0.9).

Because intergenic cheRNA (**icheRNA)**, which are defined as cheRNA without no coordinate overlap with known coding genes, is similar to eRNA [4, 5], we examined the occupancy of enhancer marks (ChIP-seq signals of EP300, H3K27ac, H3K4me1) and a repressive mark (H3K27me3) around the TSS of the 2.0 k to 6.7 k icheRNA identified by each pipeline (Fig 3c). In this analysis, we used ChromHMM [20]predicted eRNA (Fig 3c, yellow) and non-enhancer RNA (Fig 3c, black) as positive and negative controls. ChromHMM-predicted eRNA is defined as intergenic RNA that overlaps (at least 1 base, same strand) with any ChromHMM predicted “strong enhancer” region and ChromHMM-predicted non-enhancer RNA is defined as transcribed RNA that has no overlap with any predicted “strong enhancer” or “weak enhancer” region. We found that the levels of enhancer marks (EP300, H3K27ac, H3K4me1) are significantly higher around TSS of icheRNA than at the TSS of ChromHMM predicted non-enhancer RNA, while the level of repressive marks is significantly lower. Moreover, among the 4 pipelines, the icheRNA identified by Tuxedo pipeline have the highest levels of H3K27ac and H3K4me1 enhancer marks around their TSS, and relatively lower levels of repressive marks. We also noticed that the levels of enhancer marks around TSS of ChromHMM-predicted eRNA are much higher than those around TSS of icheRNA. Considering that ChromHMM predicts enhancer regions based on histone modification patterns, the enhancers predicted by ChromHMM may be biased toward having high occupancy of these canonical enhancer marks. Additionally, all three new pipelines slightly improved the cell-type specificity compared to Werner, as evaluated by the proportion of tissue-specific icheRNA identified by each pipeline (represented by R1 score in Fig 3d).

Overall, we conclude that Tuxedo and Concatenating outperform the other two pipelines in identifying expected cheRNA, and that the Tuxedo predicted icheRNA transcripts are more highly enriched in enhancer hallmarks compared to other methods. In this sense, Tuxedo outperforms Concatenating and other pipelines in enriching enhancer hallmarks in the same cell type.

### 2.3. Intergenic cheRNAs (icheRNA) uniquely present eRNA features

#### icheRNA represents a subset of noncoding RNAs de novo

Werner *et al* proposed that icheRNA is a distinct subclass of unannotated eRNA. To further examine this hypothesis, we categorized the 14k expressed nuclear RNAs detected by Tuxedo into three groups (intergenic RNA, coding-antisense RNA (labeled as “antisense RNA” in Fig 4a) and those that overlap mRNAs in the sense orientation (labeled as “mRNA” in Fig 4a). **Fig S3** shows the workflow used to categorize the three RNA groups). A large fraction (66%) of the 5,680 identified cheRNAs are transcribed from noncoding regions (Fig 4a, pink bar). In contrast, approximate 90% of the identified 5,672 sneRNAs were mRNAs (Fig 4a, blue bar). Additionally, icheRNA exhibits lower coding potential (cumulative CPC2 score [21]) than coding genes, intergenic sneRNA (isneRNA), and intergenic RNA transcribed from ChromHMM- or FANTOM-[22] predicted enhancer regions in the same cell lines (Fig 4b). The coding potential of icheRNA is therefore more similar to that of ChromHMM predicted eRNA, while that of isneRNA is more similar to that of mRNA.

**Fig 4.**
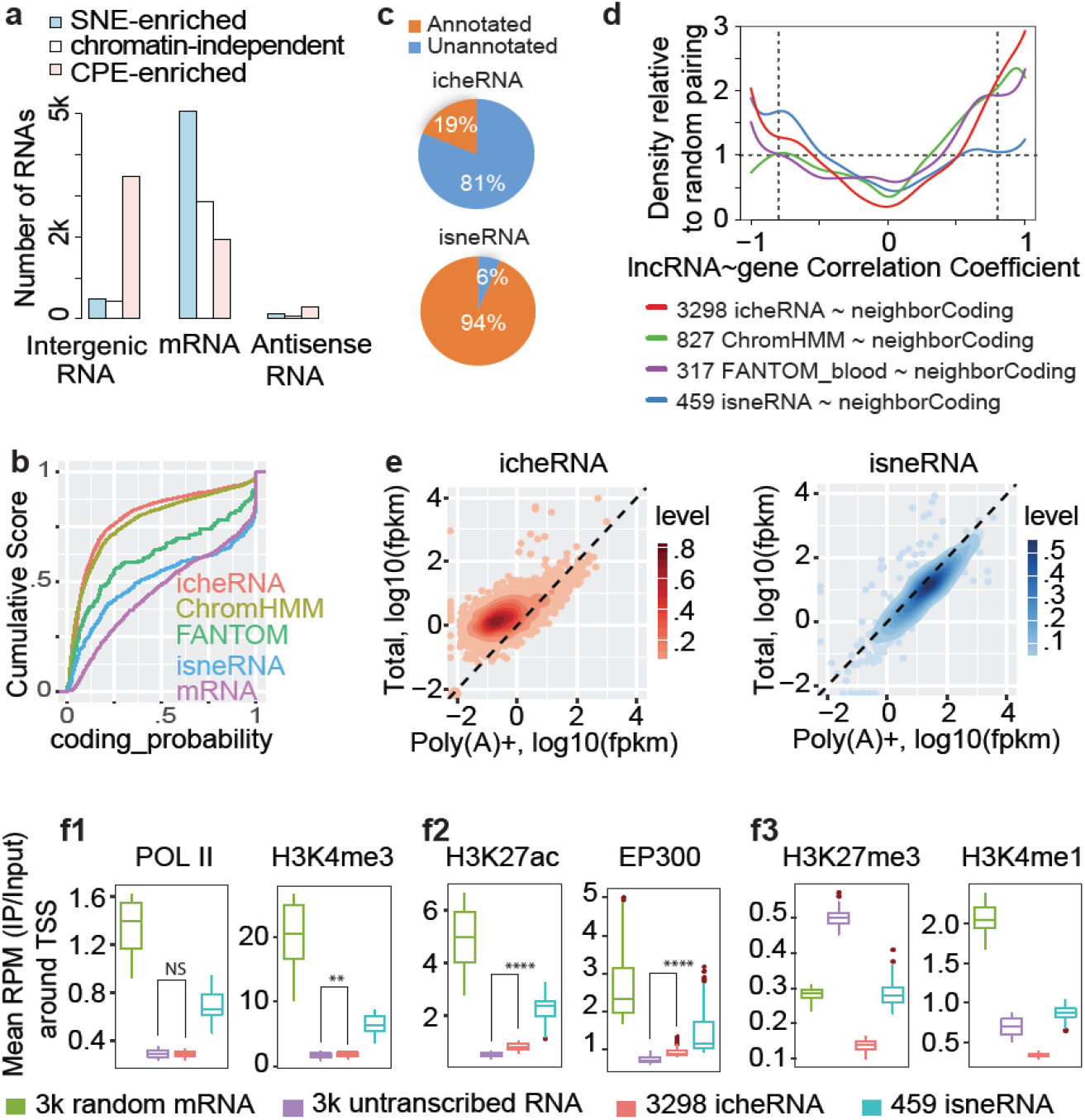
Known genomic features of the icheRNA in the K562 nucleus. **a**, Distribution of RNA classes in fractionated libraries. Three classes were defined based on their relative genomic locations to GENCODE (v25)-annotated coding genes (**Fig S3**). Chromatin-independent RNAs refer to RNAs not differentially expressed in CPE and SNE samples. **b**, Coding potential of icheRNA (red), ChromHMM predicted eRNAs (yellow), FANTOM predicted eRNAs (green), isneRNA (blue) and mRNAs (purple). Intergenic RNA overlapped (at least 1 base) with any ChromHMM/FANTOM identified enhancer region is assumed to be predicted ChromHMM/FANTOM predicted eRNAs. The online tool CPC2 is used. **c**, Percentage of GENCODE (v25) annotated and unannotated RNAs in icheRNA and isneRNA. **d**, Pairwise Correlation of expression of RNA classes. To pair an intergenic genomic feature with its neighboring gene, the adjacent upstream or downstream gene with the highest magnitude PCC is selected. The relative density at a certain PCC value is calculated by dividing the kernel density estimates of indicated RNA and neighboring coding gene pairs by that of indicated RNA and randomly selected coding gene pairs. (Two vertical dashed lines mark significant cutoffs of PCC values at −0.8 or 0.8). **e**, Normalized expression values of fractionate RNA classes. Values are given in FPKM in Poly(A)+ nuclear RNA-Seq library (x-axis, GSE88339) versus nuclear total-RNA-Seq library (y-axis, GSE87982). **f**, Average ChIP-seq read density versus input centered at promoters (±1kb centered at TSS) of RNAs, *p*-values calculated by two-sided Wilcoxon rank sum test, NS p>0.05, * p<0.01, ** p<1e-10, **** p<2.2e-16. (Note that in each panel, boxes without overlaps are significantly different without showing **** for simplicity.) 3k random mRNAs were selected from 9.8k transcribed mRNAs and 3k silent RNAs from 66.9k annotated but untranscribed RNAs.

81% (2.7 k) of the identified 3.3 k icheRNAs are previously unannotated transcripts, in contrast to only 6% (27) of the 459 isneRNAs, (Fig 4c). Additionally, over half (69% of 445) of the antisense cheRNAs are unannotated, in contrast to only 2% of 163 antisense sneRNAs. This biased annotation of noncoding RNA suggests that previously detected noncoding RNAs primarily correspond to chromatin-depleted noncoding RNA (noncoding sneRNA), and that identifying chromatin enriched RNAs from nuclear extracts can give a more balanced picture of the overall noncoding RNA population.

#### icheRNA positively correlate with adjacent genes in expression

RT-PCR experiments have shown that several eRNAs are intergenic chromatin enriched RNAs (**icheRNA**) [7]. Werner *et al.* also showed that protein-coding genes proximal to icheRNA have higher expression levels than those near to other expressed lncRNA, suggesting that icheRNA could predict cis-gene transcription [4, 5]. However, it is not clear from previous work whether higher icheRNA expression is correlated with expression of proximal protein-coding genes. To quantitatively confirm the cis-regulatory potential of icheRNA, we calculated the Pearson correlation coefficient between the expression of icheRNA and neighboring protein-coding genes, and compared it to the correlation coefficient between the expression levels of icheRNA and randomly selected protein-coding genes. isneRNA and neighboring protein-coding gene, ChromHMM predicted eRNA and neighboring protein-coding gene, and FANTOM predicted eRNA and neighboring protein-coding gene.

We find that icheRNA are more positively correlated with neighboring genes than with randomly selected genes (Fig 4d, red line shows relative density > 1 when correlation coefficient > 0.5). The same calculation for FAMTOM- or ChromHMM-predicted eRNAs, which are believed to have cis-regulatory enhancer effects, and adjacent genes in the same cell types, shows similar but weaker positive correlations. In contrast, pairs of intergenic sneRNAs (**isneRNA**) and neighboring genes (blue line) showed negative correlation (Fig 4d, blue line shows relative density > 1 when correlation coefficient < −0.5). Specifically, with a significance cutoff of correlation coefficient=0.8, 23% of the identified icheRNA transcripts, in contrast to only 11% of the isneRNA are positively correlated with proximal genes. This observation, for the first time, gives quantitative evidence for a potential cis-regulatory effect of icheRNA on adjacent genes. It also suggests that identification of icheRNA can be used as another approach to predict eRNA, comparable to approaches using ChromHMM and FANTOM database.

Transcriptional correlation analysis also displayed high relative density at correlation coefficient < −0.5 for pairs of icheRNA and neighboring coding genes (Fig 4d), indicating that not all icheRNAs are positively correlated with proximal protein-coding gene expression. Indeed, *XIST* is a canonical icheRNA that has a well-known repressive regulatory role, and it might be one of the icheRNAs that are negatively correlated with proximal protein-coding genes.

#### Polyadenylated RNA is relatively depleted in icheRNA

Most eRNAs have been reported to be unspliced and non-polyadenylated [23–26]. To test if icheRNA are similar in this regard, we compared the relative expression (measured as Reads Per Kilobase of transcript per Million mapped reads, RPKM) of intergenic cheRNAs in nuclear Poly(A)+ RNA-seq library and nuclear total RNA-seq libraries using published datasets for K562 cells (**Table S1**). We observe (Fig 4e) lower relative abundance of icheRNA in the nuclear total-RNA-seq library than in the nuclear Poly(A)+ RNA-seq library, indicating the majority of icheRNA lack polyadenylation. A similar but weaker preference for the total-RNA-seq library was also observed for antisense cheRNAs (**Fig S4**). In contrast, all protein-coding mRNAs have equivalent expression levels in two libraries, which is consistent with the role of polyadenylation in producing mRNA in the eukaryotic cell nucleus [27]. Chromatin depleted non-coding RNAs (isneRNA and antisense sneRNAs) also have similar expression levels in the two RNA-seq libraries as those of mRNAs (**Fig S4**). The patterns of polyadenylation indicate that icheRNA and isneRNA are differentially polyadenylated. With respect to polyadenylation, icheRNA is more similar to eRNA than is sneRNA, since the majority of icheRNA are not polyadenylated.

#### icheRNA and isneRNA confer different chromatin characteristics

Histone 3 lysine 4 monomethylation (H3K4me1) and histone 3 lysine 27 acetylation (H3K27ac) have been identified as key histone modification features that mark enhancers. H3K4me1 is present at both poised and active enhancers [28], while H3K27ac uniquely marks active enhancers [29]. Werner *et al*. previously observed peaks of H3K27ac near the transcriptional start sites (**TSS**) of icheRNA, however, unlike prototypical eRNA, these regions did not show abundant H3K4me1 modification [4]. To further investigate whether icheRNA have a distinct chromatin signature, we profiled the relative reads per million (RPM) of RNA polymerase II (POLII), H3K27ac, H3K4me3, H3K4me1, and H3K27me3 marks on the flanking 1 kb sequences around TSS of icheRNA, isneRNA, mRNA and unexpressed mRNA (RNAs annotated in GENCODE(v25) but not transcribed in K562) (Fig 4f). IcheRNA shows low levels of marks associated with active transcription (POLII and H3K4me3), similar to the levels of unexpressed mRNA, and lower than those of isneRNA and mRNA (Fig 4f1, red and purple box). In contrast to unexpressed mRNA, icheRNA TSS flanking regions show low levels of repressive (H3K27me3) and poised enhancer (H3K4me1) marks (Fig 4f2, red and purple box), but are enriched in active enhancer (H3K27ac and EP300) marks (Fig 4f3, red and purple box). Note that in addition to being enriched at enhancer regions, H3K27ac and EP300 are also pervasively found near TSS of actively transcribed regions. icheRNA TSS thus have a chromatin profile that is distinctly different from those of mRNA, isneRNA, and unexpressed mRNA, suggesting that significantly different modes of regulation may be controlling icheRNA expression.

In summary, icheRNA and isneRNA differ in many respects. In addition to the enrichment of specific epigenetic marks near the TSS, icheRNA has lower coding probability, lacks polyadenylation, and its expression is more positively correlated with that of neighboring coding genes. Overall icheRNA is more similar to eRNA, while isneRNA is more similar to mRNA. The similarity of icheRNA to eRNA, as defined by ChromHMM and FANTOM predictions, suggests that icheRNA identification may provide a useful independent approach to predicting eRNA.

### 2.4. Cis-regulatory potential of both icheRNA coincident with histone mark H3K9me3 and the cheRNA antisense to a coding gene

#### icheRNA with H3K9me3 signal across transcript bodies is more prone to present active cis-regulation

Histone 3 lysine 9 trimethylation (H3K9me3) is associated with constitutive heterochromatin, and has been shown to mark transcriptionally repressed regions that are mutually exclusive with H3K27me3 marked repressive regions [30–33]. We find that the levels of H3K9me3 near (100 base away from) actively transcribed icheRNA and mRNA TSS (Fig 5a1, red line and green line) are much higher than near transcriptionally silenced regions (DNA regions near to unexpressed mRNA) (Fig 5a1, purple line). In addition, H3K9me3 profiles at actively transcribed regions are quite different from those at transcriptionally silent regions: H3K9me3 signal drops at the TSS of transcribed RNA (icheRNA, isneRNA, and mRNA) (Fig 5a1, red line, blue line and green line) but is flat around TSS of unexpressed mRNA (Fig 5a1, purple line). It has been suggested, for coding transcripts, that H3K9me3 at the promoter is repressive, whereas H3K9me3 across the mRNA transcript body is activatory [31]. The pattern we observe is similar, and when combined with the previous observation that high levels of H3K9me3 modification are present in some active genes [34], it suggests that high H3K9me3 levels do not necessarily indicate transcriptional repression; H3K9me3 modification at the TSS region is more strongly associated with transcriptional silencing, in contrast, H3K9me3 at gene body regions can be actively transcribed.

**Fig 5.**
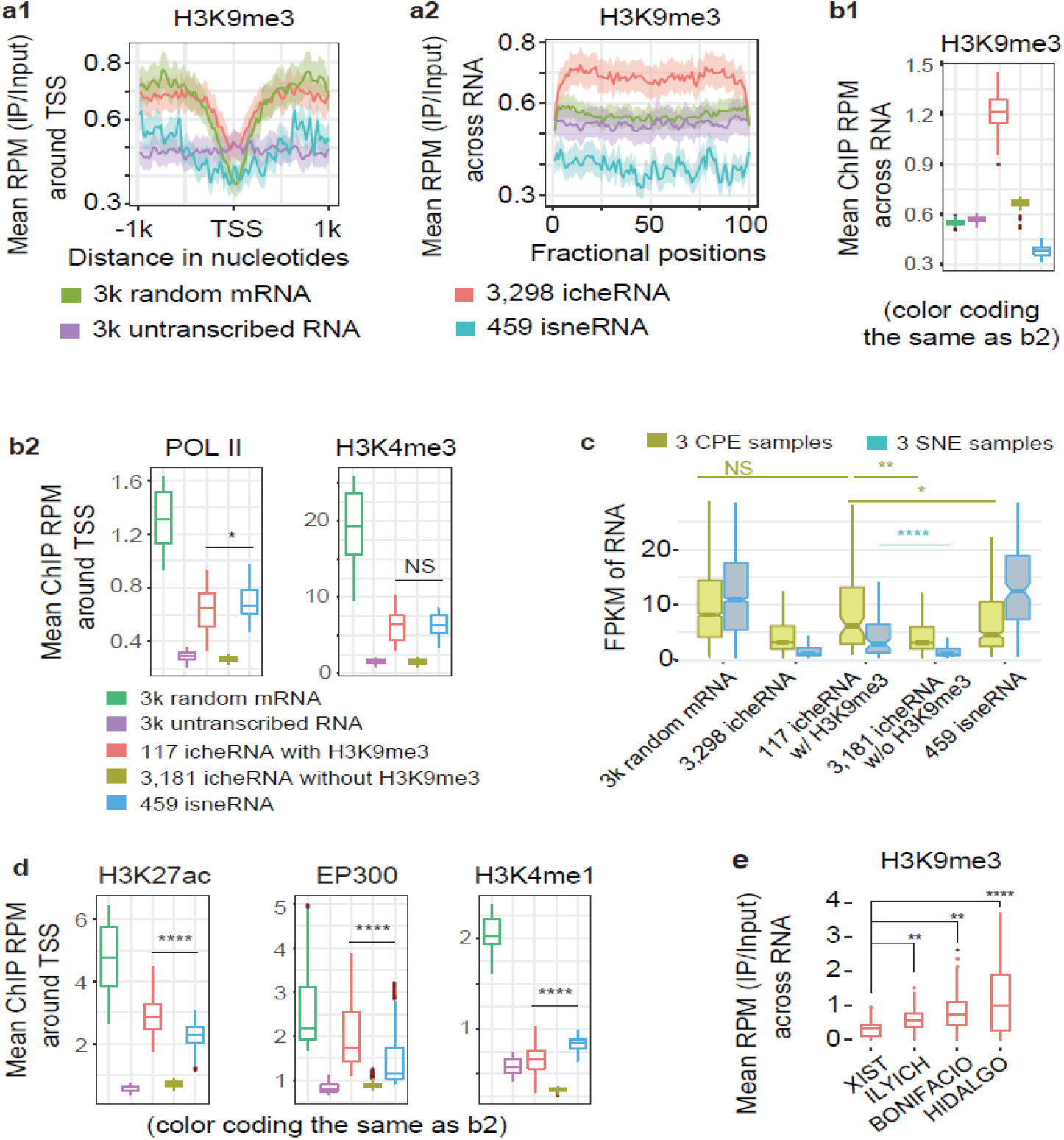
icheRNA with H3K9me3 signal concurs chromatin modification patterns of active enhancers. **a**, Average H3K9me3 ChIP-seq read density versus input in K562 cells (**a1**) at promoters (±1kb centered at TSS) of, or (**a2**) across transcript bodies. **b**, Average ChIP-seq read density versus input in K562 cells of (**b1**) H3K9me3 profiles across regions transcribing, or (**b2**) POL II and H3K4me3 profiles at promoters (±1kb centered at TSS) of five classes of RNAs (decoded in colors). **c**, Normalized expression values in FPKM in chromatin pallet extract (CPE, red boxes) and soluble nuclear extract (SNE, blue) of K562 cells for randomly selected mRNA, icheRNA, icheRNA with H3K9me3 (icheRNA w/ H3K9me3), icheRNA without H3K9me3 (icheRNA w/o H3K9me3) and isneRNA. **d**, Average ChIP-seq read density in K562 cells of active enhancer marks (H3K27ac and EP300) and poised enhancer mark (H3K4me1) profiles at promoters (±1kb centered at TSS) of five classes of RNAs. **e**, Average H3K9me3 ChIP-seq read density versus input in K562 cells across regions transcribing four canonical cheRNAs. The four cheRNAs were ordered according to their known transcriptomic regulatory functions, from the repressor (*XIST*) on the left to other three cis-activators on the right. p-values calculated by two-sided Wilcoxon rank sum test, NS p>0.05, * p<0.01, ** p<1e-10, **** p<2.2e-16.

We also note that H3K9me3 levels within the DNA region of transcribed icheRNA is substantially higher than near other transcribed RNA (*e.g.*, mRNA and isneRNA) (Fig 5a2). To further investigate the effect of H3K9me3 on icheRNA transcription, we separated DNA regions transcribing icheRNA into high H3K9me3 (at least one peak of H3K9me3 mark near the transcribed icheRNA) and low H3K9me3 (no H3K9me3 mark near the transcribed icheRNA) groups. These groups are labeled as “icheRNA with H3K9me3” and “icheRNA without H3K9me3”, respectively in Fig 5b. DNA regions in the “icheRNA with H3K9me3” have significantly higher levels of H3K9me3 modification than those in the “icheRNA without H3K9me3” group (Fig 5b1). Furthermore, chromatin signatures associated with active transcription (POL2, H3K4me3) (Fig 5b2), as well as transcription levels in both CPE and SNE samples (Fig 5c), are strikingly elevated in the “icheRNA with H3K9me3” group compared to the “icheRNA without H3K9me3” group, indicating that icheRNA are more actively transcribed within regions with high H3K9me3 modification. It also reinforces the evidence indicating that regions with abundant H3K9me3 modification can be actively transcribed.

Our previous analysis showed that icheRNA possesses features similar to eRNA, however, the TSS of icheRNA show only moderately higher levels of enhancer marks compared to unexpressed mRNA, and lower levels than TSS of isneRNA. We measured the levels of enhancer marks (H3K27ac, EP300 and H3K4me1) around TSS of “icheRNA with H3K9me3” (Fig 5d, red box). We found that levels of active enhancer marks (H3K27ac and EP300) around TSS of “icheRNA with H3K9me3” are significantly higher than at the TSS of “icheRNA without H3K9me3” (Fig 5d, yellow box) and TSS of isneRNA (Fig 5d, purple box), indicating that icheRNA surrounded by H3K9me3 marks shows high levels of active enhancer marks near the TSS, but all icheRNA do not. Moreover, we measured the H3K9me3 levels across canonical icheRNA transcribed regions and found that three previously identified icheRNA (*HIDALGO*, *ILYICH*, *BONIFACIO*) with validated positive activator functions show relatively higher H3K9me3 levels than the only icheRNA with a known repressive role (*XIST*) (Fig 5e). These examples reinforce the hypothesis that DNA regions transcribing icheRNA, even with high levels of H3K9me3 modification, may act as enhancers.

#### Antisense cheRNA (as-cheRNA) co-occurs with local mRNA silencing (Fig 6)

Antisense RNA (**asRNA**) complementary to protein-coding transcript(s) has been shown to interfere with transcription of mRNA on the opposite strand [35]. Consistent with this, asRNA accumulates preferentially in the nucleus associating with chromatin, we observe that almost (59%) of the identified 756 asRNAs in K562 cell nucleus are chromatin enriched and only 22% are chromatin depleted (Fig 4a), indicating a significant enrichment of asRNA in the chromatin pellet (Fig 6a). Moreover, we notice that about one third of the chromatin enriched asRNAs (antisense cheRNA, **as-cheRNA**) are unannotated while almost all chromatin depleted asRNAs (antisense sneRNA, **as-sneRNA**) are annotated (Fig 6b), suggesting that many as-cheRNA are completely novel.

**Fig 6.**
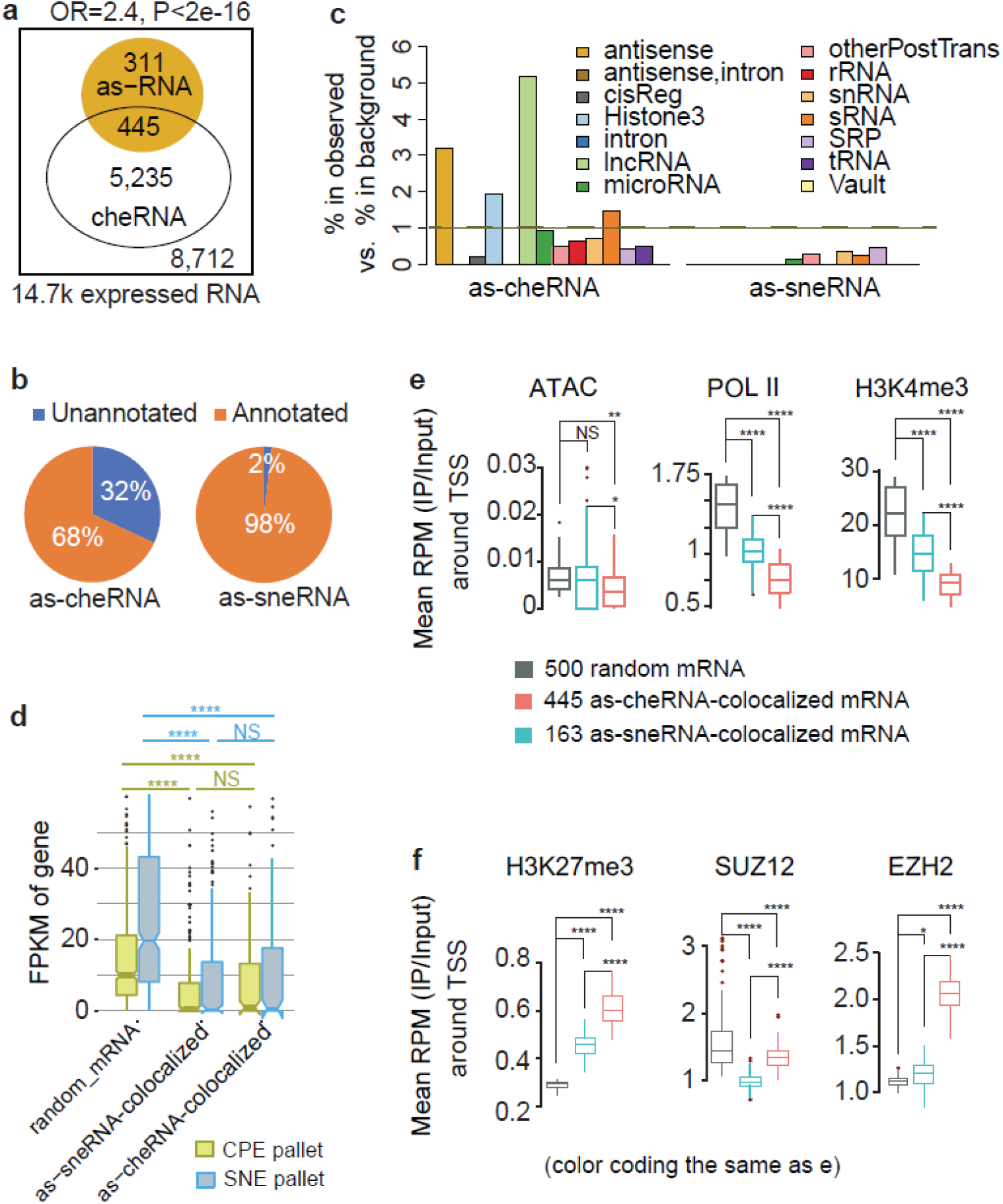
Antisense cheRNAs indicate local mRNA silencing. **a**, Venn diagram showing the enrichment of cheRNA among asRNA. Fisher’s exact test is used to estimate the strength of enrichment. Odds ration larger than 1 and p-value less than 0.05 indicate significant enrichment. **b**, Percentage of GENCODE (v25) annotated (orange) and unannotated (blue) RNA in as-cheRNA and as-sneRNA. **c**, Enrichment of 14 Rfam ncRNA secondary structure families among as-cheRNA (left sub-panel) and sneRNAs (right sub-panel). An RR-score larger than 1 indicates that as-cheRNA/as-sneRNA is overrepresented in the selected Rfam family. **d**, Normalized expression values in FPKM in chromatin pallet extract (CPE, yellow) and soluble nuclear extract (SNE, blue) of K562 cells for three RNA cleasses. **e**, Average ATAC-Seq read density and ChIP-seq read density of histone marks representing active transcription (POLII and H3K4me3) versus input in K562 cells at promoters (±1kb centered at TSS) of three RNA classes. **f**, Average ChIP-seq read density of repressive histone mark (H3K27me3) and two PRC2 subunits (SUZ12 and EZH2) versus input in K562 cells at promoters (±1kb centered at TSS) of three RNA classes. P-values are calculated using a two-sided Wilcoxon rank sum test, NS p>0.05, * p<0.01, ** p<1e-10, **** p<2.2e-16.

Regulatory mechanisms involving asRNA range from simple transcriptional interference through competing for RNA Pol II [36] to regulation of epigenomic modifications [37, 38]. A current hypothesis suggests that asRNA is acts in gene regulation at the chromatin level by recruiting epigenetic regulators, *e.g.*, polycomb repressive complex 2 (**PRC2**), to its corresponding sense mRNA to induce histone methylation and gene repression [39, 40]. Inspired by this hypothesis, we investigated a similar potential function for both as-cheRNA and as-sneRNA. Functional RNA molecules often exhibit secondary structures that are better conserved than their sequences [41], we first interrogated the sequence based predicted secondary structure of as-cheRNA and as-sneRNA in comparison to known RNA families in Rfam. Rfam collects multiple-sequence alignment-based families of RNA secondary structural motifs [41]. The motif sizes are generally less than 400 bases long **Fig S5b**), much shorter than the asRNA in the assembled transcriptome. We annotated each asRNA as belonging to a Rfam family if it fully covered a Rfam family motif. We then calculated, for each Rfam family, a) the fraction of as-cheRNA/as-sneRNA annotated to this Rfam family (the fraction in observation); b) the fraction of all assembled RNA annotated to this Rfam family (the fraction in background); and c) the ratio of the fraction in observation over the fraction background (RR-score). An RR-score larger than 1 indicates that as-cheRNA/as-sneRNA is overrepresented in the selected Rfam family. Among fourteen major RNA structural groups in the Rfam database (v13, hg38) (**Fig S5a**), three structural groups (Histone 3, lncRNA, and antisense) are significantly overrepresented in as-cheRNA (Fig 6c, two or more folds). In particular, the overrepresentation of the antisense structure group among as-cheRNA suggests that the function of as-cheRNA, rather than that of as-sneRNA, is likely to be structure-based.

We then measured the transcription level in CPE and SNE of mRNA that antisense overlaps with as-cheRNA and as-sneRNA. We find that the transcription of both as-cheRNA-colocalized mRNA and as-sneRNA-colocalized mRNA are relatively low compared to that of random mRNA (Fig 6d), suggesting a negative correlation between the transcription of sense and antisense RNA. Even though both as-cheRNA-colocalized mRNA and as-sneRNA-colocalized mRNA are shown to be repressed at similar levels, the chromatin features and histone patterns around the TSS of the two mRNA groups are significantly different. The TSS of as-cheRNA-colocalized mRNA (Fig 6e-f, red box) are significantly less open (measured by Encode ATAC-seq signal), have fewer active transcription marks (POL2, H3K4me3), but have more repressive marks (H3K27me3) and show higher PRC2 complex binding (SUZ12, EZH2) compared with random mRNA (Fig 6e-f, black box). This pattern was not observed in as-sneRNA-colocalized mRNAs (Fig 6e-f, blue box). Altogether, this suggests that as-cheRNA and as-sneRNA may cis-repress gene transcription through different mechanisms. As-cheRNA may be cis-regulatory elements that repress transcription of colocalized mRNAs on the opposite strand via recruiting the PRC2 complex to specific genomic loci.

## 3. Discussion

Detail analysis of nuclear RNA-seq sheds new light on cis-regulatory elements from lncRNA that are shorter than 1,000 bases or lowly transcribed previously embedded in noise. Operationally, cheRNA is defined by its statistically significant enrichment in chromatin after biochemical fractionation of nuclei. With our improved computational strategy, we have examined the molecular characteristics of cheRNAs in greater detail than has heretofore been possible. We find that, first, cheRNAs are more likely to be transcribed from noncoding regions, while sneRNAs are mostly transcribed from protein-coding regions. Second, icheRNA has a lower transcription level and is largely unannotated, in contrast to isneRNA which is more highly transcribed and better annotated. Traditional transcriptome profiling of non-coding RNA, using techniques such as total RNA-seq, yields the broadest survey of transcripts, but has limited ability to detect low expression transcripts such as those of icheRNA. Thus, previous analyses of noncoding RNA primarily focused on noncoding RNA with relatively high transcription levels (*e.g.*, isneRNA and as-sneRNA). In contrast, sequencing and identifying chromatin enriched RNAs in a nuclear extract more sensitively identifies low expression noncoding RNAs that previously have been ignored by conventional sequencing and analysis methods. Third, we have shown that icheRNA, in contrast to isneRNA, is mostly non-coding, non-polyadenylated, and positively correlated with the expression of neighboring coding genes (Fig 5a-5e). Notwithstanding the similarity of these features to those of eRNA, icheRNA has several unique molecular characteristics that distinguish it. For example, icheRNA is generally longer than eRNA (median length of icheRNA is ~4,400 bases; eRNA is ~350 bases, [42]) and icheRNA shows only modest coincidence with enhancer marks (H3K27ac, H3K4me1 and EP300) that are used to canonically define eRNA (Fig 4f). Moreover, some canonical icheRNA (*e.g.*, *XIST*) are known to be repressive regulators rather than activators as is eRNA. Combining all this evidence, we conclude that icheRNA and eRNAs are two distinct non-coding RNA groups that overlap. Despite there are differences, icheRNA still shows more apparent similarity to eRNAs than other RNA groups.

IcheRNA transcripts mapped to regions with high levels of H3K9me3-marks in the transcript body are more actively transcribed, and more highly associated with elevated levels of enhancer marks, than icheRNA transcribed from regions with low levels of H3K9me3 marks. This observation indicates that H3K9me3 may cover potential enhancer regions. The association between H3K9me3 and enhancers was also previously suggested by [43], who described the widespread presence of H3K9me3 at enhancer flanking regions. They also showed anecdotal examples in which regulating H3K9me3 levels at the enhancers of Mdc and Il12b affected Mdc and Il12b transcription in dendritic cells and macrophages, suggesting that H3K9me3 plays an important role in regulating enhancer activity [43]. If it can be verified that the regulatory role of H3K9me3 is a common feature of enhancers, icheRNA coincident with H3K9me3 marks may prove a very effective predictor for chromatin-based eRNA, and may be a powerful approach to predicting novel enhancer regions.

Both icheRNA and transcribed mRNA show a severe drop of H3K9me3 signal at the TSS; however, the degree to which H3K9me3 signal is decreased at the TSS is much greater for mRNA than icheRNA (Fig 5a1). As H3K9me3 is associated with condensed chromatin regions, we hypothesized that the body of icheRNAs are located within condensed chromatin regions. Agreeing with this hypothesis is the high signal of H3K9me3 along icheRNA transcript body (Fig 5a2). In metazoan cell nuclei, hundreds of large chromatin domains, termed Lamina-Associated Domains (**LADs**), are enriched in histone modification of H3K9me2 and H3K9me3, modifications that are typical of heterochromatin [44]. A study on a 1 Mb LAD encompassing the human HBB loci showed that knockdown of H3K9me3 by depletion of the two H3K9me3 methyltransferases Suv39H1 and Suv39H2 caused detachment of the LADs and nuclear lamina [45]. Considering H3K9me3 is a chromatin mark associated with closed/repressed chromatin, lncRNA transcription from H3K9me3-enriched LADs is expected to be repressed. However, the unexpected association between icheRNA and high levels of H3K9me3 chromatin marks suggests that icheRNAs may be embedded in, and actively transcribed from, condensed LADs. Indeed, we find that 48% of icheRNAs are transcribed from LADs (greater than chance expectation, empirical p < 2.2e-16), in contrast, only 12% of other RNAs are transcribed from LADs. Moreover, agreeing with the previous hypothesis by Werner *et al*. that transposable element may provide an evolutionary origin to chromatin enriched RNAs [5], we noticed that 82% of icheRNAs and 96% of icheRNAs with H3K9me3 chromatin marks in K562 overlap with class 1 transposable elements (**TEs**). Together, these observations suggest that icheRNA may represent a group of RNAs transcribed from condensed chromatin domains derived from mobile elements [13], and that the transcription of these domains is regulated in a cell-specific way.

Antisense RNA (asRNA) is another class of noncoding RNA that has been shown to have cis-regulatory functions. Consistent with previous knowledge, our analysis confirms that asRNA is more abundant in the nuclear chromatin enriched pellet than in soluble nuclear pellet. Similar to isneRNA and icheRNA, almost all as-sneRNAs are annotated, while a large fraction of as-cheRNAs lack annotation. This further suggests that sequencing RNAs abundant in the nuclear chromatin pellet can identify many novel noncoding RNAs. Despite the fact that both as-cheRNA and as-sneRNA show negative correlations in transcription level with their corresponding sense mRNA, the chromatin pattern around the TSS of as-cheRNA-colocalized mRNA and as-sneRNA-colocalized mRNA are quite different. Regions around the TSS of as-cheRNA-colocalized mRNA are less open and lack active transcription marks, but have high a level of H3K27me3 and PRC2 binding, suggesting that as-cheRNA may regulate sense mRNA transcription in cis-acting as a guide RNA for regulatory complexes that modify the target chromatin. Even though this investigation of as-cheRNA is still preliminary, it provides some testable hypotheses for asRNA function.

Chromatin-enriched nuclear RNA provides a powerful way to profile the nuclear transcriptional landscape, especially to profile the noncoding transcriptome. The computational pipeline presented here provides researchers with a reliable approach to identifying cheRNA, and for studying cell-type specific gene regulators. Although the cheRNA is unlikely to be monolithic in function, icheRNA, including icheRNA with high levels of H3K9me3 marks, may act as a transcriptional cis-activator similar to eRNA. In contrast, as-cheRNA may interact with diverse chromatin modulators to cis-repress transcription. With the Tuxedo pipeline, the future challenge will be refining the functional mechanisms of these noncoding RNA classes through exploring their regulatory roles, which are involved in diverse molecular and cellular processes in human and other organisms.

## 3. Methods

**Tables S1-2** list all publically available datasets analyzed in this study.

We compared three pipelines with the original cheRNA-identification pipeline [4]. Each pipeline includes four analytic steps: sequence mapping, transcript assembly, transcriptome construction, and signature identification (Fig 1c). Computational strategies in the latter three steps varied in four different pipelines (**S Method**). Source file for the Tuxedo pipeline is provided in the Supplementary Materials.

### Relative density of correlation between intergenic RNAs and neighbor coding genes (Fig 4d)

We calculated the pairwise Pearson correlation coefficient (**PCC**) between the intergenic RNA and protein-coding gene. We tested five types of intergenic RNA-gene groups: the icheRNA with random protein-coding gene pairs; the icheRNA with neighbor protein-coding gene pairs (icheRNA:neighborCoding); ChromHMM-predicted eRNAs with neighbor protein-coding genes pairs (ChromHMM-neighborCoding), FANTOM-predicted eRNAs with neighbor protein-coding genes pairs (FANTOM-neighborCoding) and the isneRNA with neighbor protein-coding gene pairs (isneRNA:neighborCoding).

The PCC of each intergenic RNA-gene pair was calculated based on their expression levels across all CPE and SNE samples of three cell types (K563, HEK293, H1-hESC). To pair an intergenic RNA with its neighbor protein-coding gene out of its nearest upstream and downstream genes on the same strand, the one with the highest absolute PCC value is selected. A significant cutoff of PCC values was set at −0.8 or 0.8, respectively for the negative or positive correlation. Kernel density is estimated for each intergenic RNA-gene pair group. Relative density for each intergenic RNA and neighboring protein-coding gene pairs group (e.g. icheRNA:neighborCoding) is calculated in the way of dividing the kernel density estimates of indicated intergenic RNA and neighboring protein-coding gene pairs group (e.g. icheRNA:neighborCoding) by the kernel density of icheRNA and randomly selected coding gene pairs group.

### RNA structural analysis (Fig 6c)

RNA structural analysis based on the Rfam annotations (v13, hg38) was cinducted for each Rfam-family, by dividing its frequency within a ncRNA set (icheRNA or isneRNA) versus the frequency in all assembled RNAs. Each Rfam family is represented by a multiple-sequence alignment, a consensus secondary structure, and a covariance model [41]; and we grouped one or more annotating families into a super-family according to their function as well proportions in the above noncoding transcriptome (**Fig S5a)**. The homologous ncRNA sequences in each super-family were generally less than 400 bp (2.6 on the log10-scale, **Fig S5b**), much shorter than the ncRNAs in the assembled transcriptome. Therefore, only when a ncRNA transcript fully covers an annotating family sequence, we annotated this ncRNA transcript with a Rfam super-family.

To assess the probabilities of a Rfam super-family (i) among a set of ncRNAs of interest (t), we calculated the ratio of ratios (RR) using Formula 1.

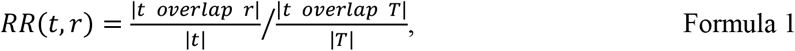

where *T* = {*t*} is the collection of all noncoding transcripts in the transcriptome, and |.| is the number of transcripts meeting a condition.

This RR score was calculated for each Rfam-family for its frequency within a ncRNA set (t) versus its global frequency. Therefore, an RR-score above 2 indicates a ncRNA set (t)-selective RNA structural family.

### Other statistics

Other statistical analyses were performed using R (**S Method**). The coding probability of RNA transcripts was calculated using Coding Potential Calculator 2 (**CPC2**) [21]. AUC was computed using the ROCR (v1.0-7) package [46].

## Acknowledgments

This work was supported by NIH R21LM012619 and through resources (BEAGLE) provided by the University of Chicago under grant 1S10OD018495-01. We would also like to acknowledge ENCODE for contributing sequencing datasets. We specifically acknowledge the assistance of Lorenzo Pesce and Purdue University ITaP Research Computing (RCAC) team for supporting high performance computing services. We thank Jeffrey Steimle and Kohta Ikegami for discussion on H3K9me3 patterns.

## Supplemental Information

Supplemental Information includes Supplementary Methods, one script, five figures, and three tables and can be found with this article online.

## Author Contributions

XY initiated this work and supervised this work together with MG. XS contributed in collecting datasets, designing pipelines, performing evaluation and analysis work, and interpreting results, with input from XY, MG, IM and AR. ZW helped with Rfam data analysis (Fig 6c), transcription factor data collection (Table S2), and transcription factor occupancy analysis (Figs 1, S1). CP participated in implicating pipelines at early stage of the project. XS wrote the manuscript, with comments from MG. XY designed pipelines, contributed to the discussion, wrote the abstract, part of introduction, and Rfam method of the manuscript, with comments from MG, IM. XS, XY revised the manuscript, with comments from MG, IM. All authors approved the final manuscript.

## Reference

1. Quinodoz S, Guttman M. Long noncoding RNAs: an emerging link between gene regulation and nuclear organization. Trends Cell Biol. 2014;24(11):651–63. doi: 10.1016/j.tcb.2014.08.009. PubMed PMID: 25441720; PubMed Central PMCID: PMCPMC4254690.

2. Khalil AM, Guttman M, Huarte M, Garber M, Raj A, Rivea Morales D, et al. Many human large intergenic noncoding RNAs associate with chromatin-modifying complexes and affect gene expression. Proc Natl Acad Sci U S A. 2009;106(28):11667–72. Epub 2009/07/03. doi: 10.1073/pnas.0904715106. PubMed PMID: 19571010; PubMed Central PMCID: PMC2704857.

3. Sun Q, Hao Q, Prasanth KV. Nuclear Long Noncoding RNAs: Key Regulators of Gene Expression. Trends Genet. 2018;34(2):142–57. doi: 10.1016/j.tig.2017.11.005. PubMed PMID: 29249332; PubMed Central PMCID: PMCPMC6002860.

4. Werner MS, Ruthenburg AJ. Nuclear Fractionation Reveals Thousands of Chromatin-Tethered Noncoding RNAs Adjacent to Active Genes. Cell Rep. 2015;12(7):1089–98. doi: 10.1016/j.celrep.2015.07.033. PubMed PMID: 26257179; PubMed Central PMCID: PMCPMC5697714.

5. Werner MS, Sullivan MA, Shah RN, Nadadur RD, Grzybowski AT, Galat V, et al. Chromatin-enriched lncRNAs can act as cell-type specific activators of proximal gene transcription. Nat Struct Mol Biol. 2017;24(7):596–603. doi: 10.1038/nsmb.3424. PubMed PMID: 28628087; PubMed Central PMCID: PMCPMC5682930.

6. Gayen S, Kalantry S. Chromatin-enriched lncRNAs: a novel class of enhancer RNAs. Nat Struct Mol Biol. 2017;24(7):556–7. doi: 10.1038/nsmb.3430. PubMed PMID: 28686230; PubMed Central PMCID: PMCPMC6190603.

7. Yang XH, Nadadur RD, Hilvering CR, Bianchi V, Werner M, Mazurek SR, et al. Transcription-factor-dependent enhancer transcription defines a gene regulatory network for cardiac rhythm. Elife. 2017;6. doi: 10.7554/eLife.31683. PubMed PMID: 29280435; PubMed Central PMCID: PMCPMC5745077.

8. Griffith M, Walker JR, Spies NC, Ainscough BJ, Griffith OL. Informatics for RNA Sequencing: A Web Resource for Analysis on the Cloud. PLoS Comput Biol. 2015;11(8):e1004393. doi: 10.1371/journal.pcbi.1004393. PubMed PMID: 26248053; PubMed Central PMCID: PMCPMC4527835.

9. Kumar A, Kankainen M, Parsons A, Kallioniemi O, Mattila P, Heckman CA. The impact of RNA sequence library construction protocols on transcriptomic profiling of leukemia. BMC Genomics. 2017;18(1):629. doi: 10.1186/s12864-017-4039-1. PubMed PMID: 28818039; PubMed Central PMCID: PMCPMC5561555.

10. Harrow J, Frankish A, Gonzalez JM, Tapanari E, Diekhans M, Kokocinski F, et al. GENCODE: the reference human genome annotation for The ENCODE Project. Genome Res. 2012;22(9):1760–74. doi: 10.1101/gr.135350.111. PubMed PMID: 22955987; PubMed Central PMCID: PMC3431492.

11. Hunter RG, Murakami G, Dewell S, Seligsohn M, Baker ME, Datson NA, et al. Acute stress and hippocampal histone H3 lysine 9 trimethylation, a retrotransposon silencing response. Proc Natl Acad Sci U S A. 2012;109(43):17657–62. doi: 10.1073/pnas.1215810109. PubMed PMID: 23043114; PubMed Central PMCID: PMCPMC3491472.

12. Noh KM, Maze I, Zhao D, Xiang B, Wenderski W, Lewis PW, et al. ATRX tolerates activity-dependent histone H3 methyl/phos switching to maintain repetitive element silencing in neurons. Proc Natl Acad Sci U S A. 2015;112(22):6820–7. doi: 10.1073/pnas.1411258112. PubMed PMID: 25538301; PubMed Central PMCID: PMCPMC4460490.

13. Xie X, Kamal M, Lander ES. A family of conserved noncoding elements derived from an ancient transposable element. Proc Natl Acad Sci U S A. 2006;103(31):11659–64. doi: 10.1073/pnas.0604768103. PubMed PMID: 16864796; PubMed Central PMCID: PMCPMC1518811.

14. Derrien T, Johnson R, Bussotti G, Tanzer A, Djebali S, Tilgner H, et al. The GENCODE v7 catalog of human long noncoding RNAs: analysis of their gene structure, evolution, and expression. Genome Res. 2012;22(9):1775–89. Epub 2012/09/08. doi: 10.1101/gr.132159.111. PubMed PMID: 22955988; PubMed Central PMCID: PMC3431493.

15. Bourgon R, Gentleman R, Huber W. Independent filtering increases detection power for high-throughput experiments. Proc Natl Acad Sci U S A. 2010;107(21):9546–51. doi: 10.1073/pnas.0914005107. PubMed PMID: 20460310; PubMed Central PMCID: PMCPMC2906865.

16. Hart T, Komori HK, LaMere S, Podshivalova K, Salomon DR. Finding the active genes in deep RNA-seq gene expression studies. BMC Genomics. 2013;14:778. doi: 10.1186/1471-2164-14-778. PubMed PMID: 24215113; PubMed Central PMCID: PMCPMC3870982.

17. Ritchie ME, Phipson B, Wu D, Hu Y, Law CW, Shi W, et al. limma powers differential expression analyses for RNA-sequencing and microarray studies. Nucleic Acids Res. 2015;43(7):e47. doi: 10.1093/nar/gkv007. PubMed PMID: 25605792; PubMed Central PMCID: PMCPMC4402510.

18. Niknafs YS, Pandian B, Iyer HK, Chinnaiyan AM, Iyer MK. TACO produces robust multisample transcriptome assemblies from RNA-seq. Nat Methods. 2017;14(1):68–70. doi: 10.1038/nmeth.4078. PubMed PMID: 27869815; PubMed Central PMCID: PMCPMC5199618.

19. Tilgner H, Knowles DG, Johnson R, Davis CA, Chakrabortty S, Djebali S, et al. Deep sequencing of subcellular RNA fractions shows splicing to be predominantly co-transcriptional in the human genome but inefficient for lncRNAs. Genome Res. 2012;22(9):1616–25. doi: 10.1101/gr.134445.111. PubMed PMID: 22955974; PubMed Central PMCID: PMCPMC3431479.

20. Ernst J, Kellis M. ChromHMM: automating chromatin-state discovery and characterization. Nat Methods. 2012;9(3):215–6. doi: 10.1038/nmeth.1906. PubMed PMID: 22373907; PubMed Central PMCID: PMCPMC3577932.

21. Kang YJ, Yang DC, Kong L, Hou M, Meng YQ, Wei L, et al. CPC2: a fast and accurate coding potential calculator based on sequence intrinsic features. Nucleic Acids Res. 2017;45(W1):W12–W6. doi: 10.1093/nar/gkx428. PubMed PMID: 28521017; PubMed Central PMCID: PMCPMC5793834.

22. de Hoon M, Shin JW, Carninci P. Paradigm shifts in genomics through the FANTOM projects. Mamm Genome. 2015;26(9-10):391–402. doi: 10.1007/s00335-015-9593-8. PubMed PMID: 26253466; PubMed Central PMCID: PMCPMC4602071.

23. Kim TK, Hemberg M, Gray JM, Costa AM, Bear DM, Wu J, et al. Widespread transcription at neuronal activity-regulated enhancers. Nature. 2010;465(7295):182–7. doi: 10.1038/nature09033. PubMed PMID: 20393465; PubMed Central PMCID: PMCPMC3020079.

24. De Santa F, Barozzi I, Mietton F, Ghisletti S, Polletti S, Tusi BK, et al. A large fraction of extragenic RNA pol II transcription sites overlap enhancers. PLoS Biol. 2010;8(5):e1000384. doi: 10.1371/journal.pbio.1000384. PubMed PMID: 20485488; PubMed Central PMCID: PMCPMC2867938.

25. Lam MT, Li W, Rosenfeld MG, Glass CK. Enhancer RNAs and regulated transcriptional programs. Trends Biochem Sci. 2014;39(4):170–82. doi: 10.1016/j.tibs.2014.02.007. PubMed PMID: 24674738; PubMed Central PMCID: PMCPMC4266492.

26. Kim TK, Hemberg M, Gray JM. Enhancer RNAs: a class of long noncoding RNAs synthesized at enhancers. Cold Spring Harb Perspect Biol. 2015;7(1):a018622. doi: 10.1101/cshperspect.a018622. PubMed PMID: 25561718; PubMed Central PMCID: PMCPMC4292161.

27. Guhaniyogi J, Brewer G. Regulation of mRNA stability in mammalian cells. Gene. 2001;265(1-2):11–23. PubMed PMID: 11255003.

28. Dorighi KM, Swigut T, Henriques T, Bhanu NV, Scruggs BS, Nady N, et al. Mll3 and Mll4 Facilitate Enhancer RNA Synthesis and Transcription from Promoters Independently of H3K4 Monomethylation. Mol Cell. 2017;66(4):568–76 e4. doi: 10.1016/j.molcel.2017.04.018. PubMed PMID: 28483418; PubMed Central PMCID: PMCPMC5662137.

29. Creyghton MP, Cheng AW, Welstead GG, Kooistra T, Carey BW, Steine EJ, et al. Histone H3K27ac separates active from poised enhancers and predicts developmental state. Proc Natl Acad Sci U S A. 2010;107(50):21931–6. Epub 2010/11/26. doi: 10.1073/pnas.1016071107. PubMed PMID: 21106759; PubMed Central PMCID: PMC3003124.

30. Hublitz P, Albert M, Peters AH. Mechanisms of transcriptional repression by histone lysine methylation. Int J Dev Biol. 2009;53(2-3):335–54. doi: 10.1387/ijdb.082717ph. PubMed PMID: 19412890.

31. Kouzarides T. Chromatin modifications and their function. Cell. 2007;128(4):693–705. doi: 10.1016/j.cell.2007.02.005. PubMed PMID: 17320507.

32. Zhang T, Cooper S, Brockdorff N. The interplay of histone modifications - writers that read. EMBO Rep. 2015;16(11):1467–81. doi: 10.15252/embr.201540945. PubMed PMID: 26474904; PubMed Central PMCID: PMCPMC4641500.

33. Becker JS, Nicetto D, Zaret KS. H3K9me3-Dependent Heterochromatin: Barrier to Cell Fate Changes. Trends Genet. 2016;32(1):29–41. doi: 10.1016/j.tig.2015.11.001. PubMed PMID: 26675384; PubMed Central PMCID: PMCPMC4698194.

34. Barski A, Cuddapah S, Cui K, Roh TY, Schones DE, Wang Z, et al. High-resolution profiling of histone methylations in the human genome. Cell. 2007;129(4):823–37. doi: 10.1016/j.cell.2007.05.009. PubMed PMID: 17512414.

35. Tufarelli C, Stanley JA, Garrick D, Sharpe JA, Ayyub H, Wood WG, et al. Transcription of antisense RNA leading to gene silencing and methylation as a novel cause of human genetic disease. Nat Genet. 2003;34(2):157–65. doi: 10.1038/ng1157. PubMed PMID: 12730694.

36. Shearwin KE, Callen BP, Egan JB. Transcriptional interference--a crash course. Trends Genet. 2005;21(6):339–45. doi: 10.1016/j.tig.2005.04.009. PubMed PMID: 15922833; PubMed Central PMCID: PMCPMC2941638.

37. Kotake Y, Nakagawa T, Kitagawa K, Suzuki S, Liu N, Kitagawa M, et al. Long non-coding RNA ANRIL is required for the PRC2 recruitment to and silencing of p15(INK4B) tumor suppressor gene. Oncogene. 2011;30(16):1956–62. doi: 10.1038/onc.2010.568. PubMed PMID: 21151178; PubMed Central PMCID: PMCPMC3230933.

38. Bhan A, Mandal SS. LncRNA HOTAIR: A master regulator of chromatin dynamics and cancer. Biochim Biophys Acta. 2015;1856(1):151–64. doi: 10.1016/j.bbcan.2015.07.001. PubMed PMID: 26208723; PubMed Central PMCID: PMCPMC4544839.

39. Magistri M, Faghihi MA, St Laurent G, 3rd, Wahlestedt C. Regulation of chromatin structure by long noncoding RNAs: focus on natural antisense transcripts. Trends Genet. 2012;28(8):389–96. doi: 10.1016/j.tig.2012.03.013. PubMed PMID: 22541732; PubMed Central PMCID: PMCPMC3768148.

40. Latge G, Poulet C, Bours V, Josse C, Jerusalem G. Natural Antisense Transcripts: Molecular Mechanisms and Implications in Breast Cancers. Int J Mol Sci. 2018;19(1). doi: 10.3390/ijms19010123. PubMed PMID: 29301303; PubMed Central PMCID: PMCPMC5796072.

41. Kalvari I, Argasinska J, Quinones-Olvera N, Nawrocki EP, Rivas E, Eddy SR, et al. Rfam 13.0: shifting to a genome-centric resource for non-coding RNA families. Nucleic Acids Res. 2018;46(D1):D335–D42. doi: 10.1093/nar/gkx1038. PubMed PMID: 29112718; PubMed Central PMCID: PMCPMC5753348.

42. Andersson R, Gebhard C, Miguel-Escalada I, Hoof I, Bornholdt J, Boyd M, et al. An atlas of active enhancers across human cell types and tissues. Nature. 2014;507(7493):455–61. doi: 10.1038/nature12787. PubMed PMID: 24670763.

43. Zhu Y, van Essen D, Saccani S. Cell-type-specific control of enhancer activity by H3K9 trimethylation. Mol Cell. 2012;46(4):408–23. doi: 10.1016/j.molcel.2012.05.011. PubMed PMID: 22633489.

44. van Steensel B, Belmont AS. Lamina-Associated Domains: Links with Chromosome Architecture, Heterochromatin, and Gene Repression. Cell. 2017;169(5):780–91. doi: 10.1016/j.cell.2017.04.022. PubMed PMID: 28525751; PubMed Central PMCID: PMCPMC5532494.

45. Bian Q, Khanna N, Alvikas J, Belmont AS. beta-Globin cis-elements determine differential nuclear targeting through epigenetic modifications. J Cell Biol. 2013;203(5):767–83. doi: 10.1083/jcb.201305027. PubMed PMID: 24297746; PubMed Central PMCID: PMCPMC3857487.

46. Sing T, Sander O, Beerenwinkel N, Lengauer T. ROCR: visualizing classifier performance in R. Bioinformatics. 2005;21(20):3940–1. doi: 10.1093/bioinformatics/bti623. PubMed PMID: 16096348.

